# Prudent burrow-site selection in a landscape of fear

**DOI:** 10.1101/2023.05.22.541844

**Authors:** Viraj R. Torsekar, Aparna Lajmi, Dror Hawlena

**Affiliations:** Risk-Management Ecology Lab, Department of Ecology, Evolution & Behavior, The Alexander Silberman Institute of Life Sciences, The Hebrew University of Jerusalem; Theoretical Ecology and Evolution Lab, Centre for Ecological Sciences, Indian Institute of Science; Institute of Evolution, Department of Evolutionary and Environmental Biology, University of Haifa

**Keywords:** phenotype-environment matching, prudent habitat choice, assortative mating, predation risk, non-consumptive effects, desert isopods

## Abstract

Prey should select safer breeding sites over riskier sites of otherwise similar habitats. This preference, however, may differ between conspecifics of different competitive abilities if the costs of intraspecific competition overpower the benefits of breeding in a safer site. Our goal was to test this hypothesis by exploring the burrow-site selection of different-sized desert isopods (*Hemilepistus reaumuri*) near and away from a scorpion burrow. We found that larger females are more likely to occupy burrows than smaller females, regardless of whether these burrows were close or away from scorpion burrows. We also found that larger females stayed longer in safer burrows and that smaller females tended to stay longer in riskier sites even in the absence of direct competition, implying a prudent burrow-site selection. We found no association between male size and the tendency to occupy or to spend time in a burrow, regardless of whether these burrows were close or away from scorpion burrows. Our work highlights the need to consider intraspecific competition when exploring how predators regulate prey behavior.

## Introduction

Nest-site selection is a key determinant of animal fitness, with profound ecological and evolutionary consequences [1]. Preferred nest-sites, among other things, should maximize food provisioning and minimize adverse consequences of climate and predation on both parents and offspring [2]. Predation may adversely affect the fitness of nesting individuals by reducing their survivorship and future reproductions [3,4], decreasing offspring survivorship, and inflicting long-lasting fitness costs in surviving offspring [5]. Thus, all else being equal, prey should select nest sites with low predation risk [6]. In a heterogeneous landscape of fear (i.e., spatiotemporal variation in predation risk [7]) a limited number of safe nest sites can generate intense intraspecific competition favoring conspecifics that can better identify, occupy and defend these sites [8]. This may lead to non-random prey distribution across riskier and safer sites [9]. Determining how predation-risk affects nest-site selection by conspecifics of different phenotypes may assist in explaining spatial distribution of different prey phenotypes across the landscape of fear.

Two mechanisms can lead to phenotype-environment matching by intraspecific competition, namely (a) aggressive interactions [10] and (b) prudent behavior by inferior competitors [11]. Similar nest-site preference by individuals of different phenotypes may lead to antagonistic interaction in which the inferior, often smaller or less aggressive individual is likely to lose and be expelled from the preferred site [12]. Alternatively, inferior competitors may exhibit ‘prudent behaviour’ where they proactively select low-quality habitats to avoid the costs of engaging in aggressive interactions, leaving high-quality habitats to better competitors that are more likely to win such encounters [11]. Also, prudent nest-site selection may allow inferior individuals to promptly ensure access to a nest site, even a risky one, rather than taking the potentially fatal chance of not finding a nest site at all. Prudent nest-site selection is, thus, expected in heterogeneous environments where competition for safer sites is high and the associated direct and indirect costs of losing a fight are severe [13,14]. Identifying whether aggressive interactions or prudent behavior governs prey nest site selection may shed new light on how predators regulate spatial divergence of prey populations.

We aimed to examine whether prudent behavior or aggressive antagonistic interactions govern the desert isopods (*Hemilepistus reaumuri*) burrow-site selection near and away from Israeli gold scorpion (*Scorpio maurus*) predators, using a manipulative field experiment in the Negev Desert, Israel. Isopod density in the Negev can reach up to 48 individuals per 1 m^2^ [15]. Isopods are monogamous and live with their approximately 70 offspring in a single permanent burrow. In February, isopods disperse from their natal burrows, form pairs and establish new family burrows. At this time, competition for burrows sites is severe as reflected by the large number of isopods that remain under plant-debris and stones after not finding a burrow. Females establish burrows often by occupying existing holes, empty burrows, or even small soil depressions that they deepen. Males choose and fight over females, and larger males are more likely to win these encounters [16]. Isopod pairs exhibit size assortative mating, and this pattern is regulated by predation risk [17]. Specifically, proximity to a scorpion burrow reduces the reproductive value of females occupying those burrows, resulting in smaller males settling in them. Isopods estimate scorpion predation risk by integrating various safety and predator cues ([18,19]. Desert isopods exhibit biparental care and the offspring remain within their parental burrow throughout their developmental stage, underscoring the importance of selecting a safe burrow site. Females have not been observed to fight over burrows [16]. Thus, we hypothesized that female isopods of larger size should select safer nest sites and female of smaller size should select riskier sites, exhibiting prudent burrow site selection. We did not form specific hypotheses for males since they do not initiate burrows but choose and fight over burrows occupied by females. Nonetheless male behavior was observed and reported as it was difficult to sex isopods without handling them, and as a reference for the observed female behavior.

## Methods

In March 2021, we conducted a field experiment at the Avdat Research Station, Negev desert, Israel (30°47’02” N, 34°46’09” E; Fig. S2C). To explore the isopods’ nest-site selection, we established 14 experimental blocks, each including two groups of four 0.8 cm diameter x 4 cm depth cylindrical holes that were dug in the corners of an imaginary 50 cm diagonal square (Fig. S2A). We drilled the holes using a 0.7 mm drill-bit. The minimal distance between the two burrow-groups was one meter. In each block, we dug a 10 cm long scorpion-burrow in the middle of one randomly chosen burrow-group (Fig. S2B). The neighboring burrow-group in each block served as a no-scorpion “safe” control. To all scorpion-burrows, we introduced scorpions of similar size. We encouraged the scorpion to occupy the hole by surrounding it overnight with small enclosures (details in [17]). We covered all holes with plastic lids immediately after digging them to prevent isopods from inspecting the holes prior to the beginning of the experiment. No blocks included natural burrows of isopods or scorpions.

Each experiment lasted for two consecutive days. At the peak of the isopods’ daily dispersal activity, between 13:00 and 14:00 hours, we removed all lids to begin the experiment. At the end of day one between 16:00 and 17:00 hours, the experiment was paused due to reduced isopod activity. All burrows were then covered with lids to ensure that animals inside the burrow would not escape and to prevent new hole occupancy. On day two, the experiment was restarted by removing the lids between 13:00 and 14:00 hours. At the end of day two, all isopods still inside burrows were collected, measured and released. For isopods that stayed inside burrows between the two days, durations across both days were disregarded from the analysis to discount the time when the experiment was not running.

During the experiment, each block was monitored by one person. Any isopod visiting a hole was focally observed. Their activity near and inside the hole was recorded including whether the isopod occupied the hole or rejected it, and if occupied, for how long did the isopod stay inside. Rejected holes were classified as either those that were not occupied regardless of how long the isopod spent near them or holes that were briefly occupied. Brief occupation was defined using a cut-off based on the bimodal nature of the duration data (Fig. S1). Isopods that came in contact with a hole were temporarily collected after departing from the experimental blocks, and then were sexed, weighed, temporarily marked and released. This procedure prevented handling-stress during the experiment, and ensured that each isopod was sampled only once. Body weight in desert isopods is a good proxy for size [17]. For the burrow occupation analyses, we considered only visits to non-occupied holes to ensure that isopod decision-making is influenced only by the habitat and not by the presence of a conspecific. Due to small sample size, we could not include the visits to occupied burrows when examining the time spent within the burrows (see S1).

We used a logistic generalized linear mixed model (GLMM) to investigate what drives an isopod’s choice to occupy or reject a hole, with predation risk, isopod body weight and their interaction as the fixed predictors. To examine what predicts how long an isopod spends inside the burrow, we ran a linear mixed model (LMM) with predation risk, isopod body weight and their interaction as the fixed predictors. Block was the random effect in all subsequent models. Since only 5 individuals interacted with more than one hole, individual identity was not considered as a nested random effect. Finally, we performed permutation tests to investigate whether more females or males occupied holes till the end of the trials. All analyses were performed using R version 4.1.2 [20], models were run using the packages “glmmTMB” [21] or “lme4” [22]. Post-hoc analyses to obtain estimated marginal means (EMMs) were performed using package “emmeans” [23].

## Results

The likelihood of female isopods occupying a burrow they visit increased with their size (Slope: χ^2^ = 4.718, Df = 1, p = 0.029; N= 37; Fig. 1A) regardless of the risk at the burrow (Intercept: χ^2^ = 1.215, Df = 1, p = 0. 27; N= 37; Fig. 1A). The decision of male isopods, however, was unaffected by either their own size (Slope: χ^2^ = 0.063, Df = 1, p = 0.802; N= 50; Fig. 1B) or proximity of scorpion to the burrow (Intercept: χ^2^ = 2.812, Df = 1, p = 0.094; N= 50; Fig. 1A). Interaction terms were dropped from the analyses since they were not significant for both females (χ^2^ = 3.155, Df = 1, p = 0.09; N= 37) and males (χ^2^ = 1.206, Df = 1, p = 0.28; N= 50).

**Figure 1.**
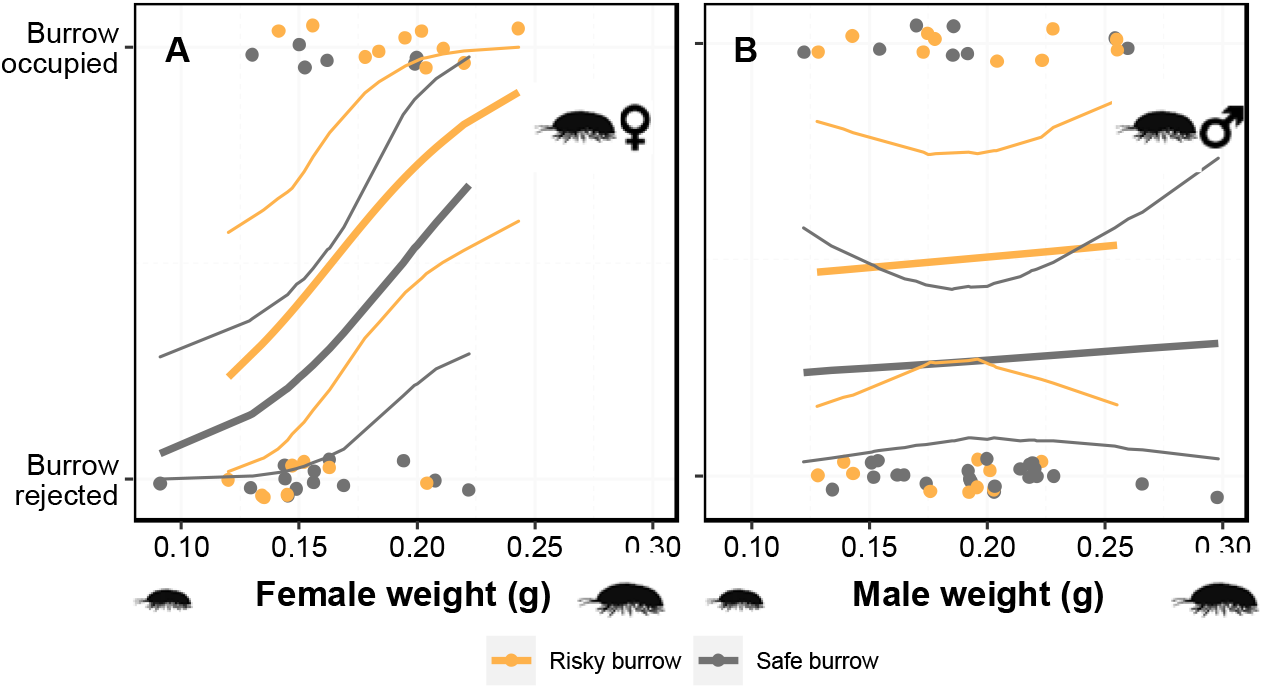
Isopod decision to reject or occupy a burrow was dependent on individual weight for (A) females but not habitat quality, (B) and neither for males. Model predictions along with 95% confidence intervals shown.

Once occupied, the time females spent inside the burrows depended on both their body size and risk at the burrow (Interaction term: χ^2^ = 13.747, Df = 1, p < 0.001; Slope: χ^2^ = 0.012, Df = 1, p = 0.91; Intercept: χ^2^ = 0.439, Df = 1, p = 0.51; N = 16; Fig. 2A). Post-hoc analyses revealed that larger females prefer safer burrows (post hoc, t-ratio = −2.768, p = 0.01), whereas risky burrows elicited an opposite trend although not significant (post hoc, t-ratio = −1.648, p = 0.13). On the other hand, the time males spent in the burrows was not influenced by either body size (Slope: χ^2^ = 0.262, Df = 1, p = 0.6; N = 18; Fig. 2B) or risk (Intercept: χ^2^ = 1.354, Df = 1, p = 0. 24; N = 18; Fig. 2B). For males, the interaction-term was not significant (χ^2^ = 0.0009, Df = 1, p = 0.97; N= 18). Thus, we did not include the interaction in the final analysis. Most individuals that occupied the experimental holes till the end of the experiment were females (permutation test, Number of simulations = 10000, p < 0.001), corroborating our understanding that males do not settle in burrows unless a female is present first.

**Figure 2.**
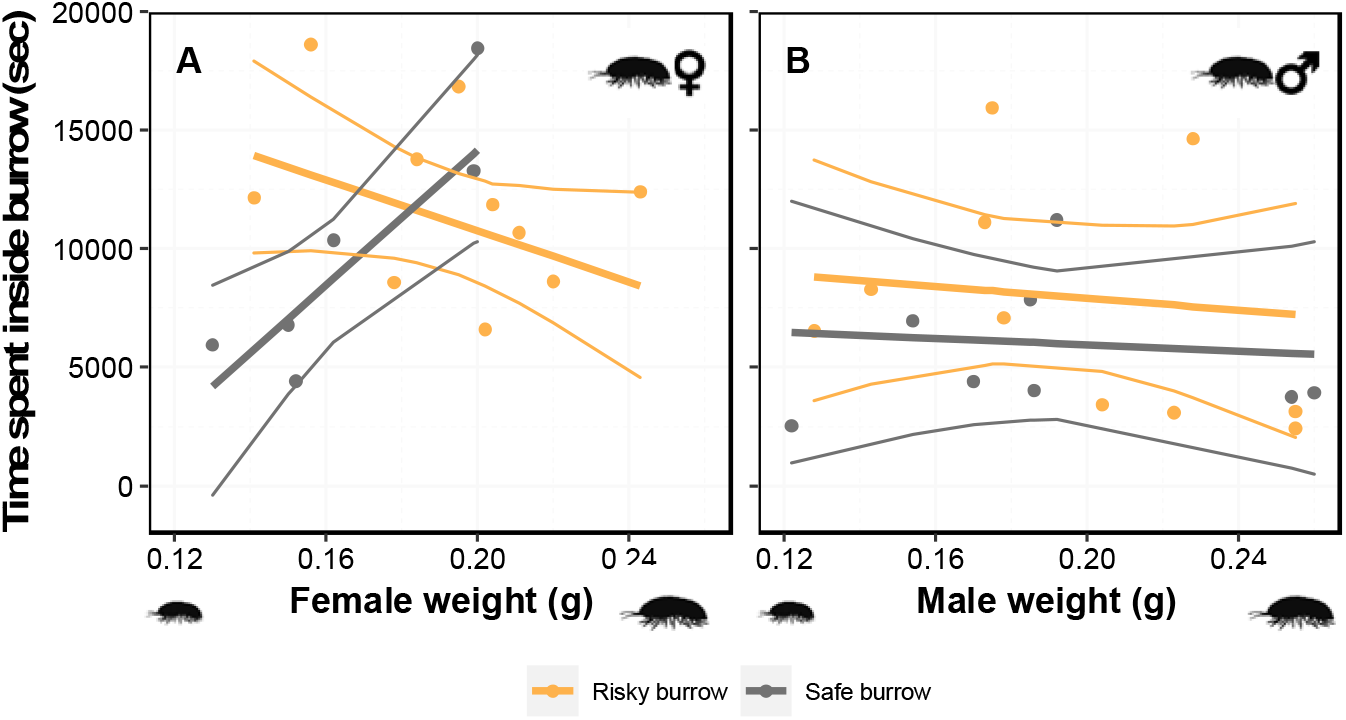
Time duration that isopods spent inside occupied burrows was dependent on both habitat quality and individual weight for (A) females (B) but neither for males. Model predictions along with 95% confidence intervals shown.

## Discussion

Using a field experiment in which dispersing isopods could select burrow sites near and away from scorpion predators, we showed that female isopods adjust their burrow-site selection based on their own body size, and that smaller females use prudent behavior instead of always favoring safer sites. Large females were more likely to enter a burrow instead of rejecting it, regardless of whether or not this burrow was near or away from a scorpion burrow. Females could identify risky sites and altered their behavior accordingly. In safe sites larger females stayed longer within burrows than smaller females, while in risky sites smaller females tended to spend more time in burrows than larger females. We found no association between male size and the tendency to occupy or to spend time in a burrow, regardless of whether these burrows were close or away from scorpion burrows.

Previous work found that time to settling is positively associated with isopods’ family success; i.e., survival until autumn [24], implying that females benefit from sampling different burrows before settling in one. This strategy seems rather risky given the high density of dispersing isopods and the limited number of potential burrow sites. We found that burrow-site selection by female isopods was regulated by body size, regardless of whether the burrow was near or away from scorpion burrows. Large females were more likely to temporarily occupy the artificial burrows than smaller females. These different burrow-selection strategies may reflect size dependent differences in females’ ability to displace resident females from already occupied burrows. Smaller females may prefer occupying a burrow that they are most likely to reside in, while lager females can take the risk of temporarily occupying a burrow and replacing it later. Thus, we suggest that smaller females adopt risk averse behavior, attempting to gather enough information that may allow them to choose an adequate burrow that they are less likely to leave, and can better defend in the future.

Large females stayed longer in safe burrows than in burrows that were close to scorpion burrows. Smaller females did not show this preference and tended to stay longer in risky burrows even in the absence of direct competition. This suggests that small females consider their self-phenotype when searching for a new burrow site, leading to a prudent burrow site selection. Theory predicts prudent choice by less-competitive individuals when the costs associated with competitive interactions are greater than the expected fitness benefits of choosing a better option [25,26]. In our system, all females should choose burrow site that are farther away from scorpion burrows to reduce predation risks on both parents and offspring. Our results suggest that the direct costs of engaging in competitive interactions and the increasing probability of future displacement by larger females surpass the benefits of living in a less risky environment.

Prudent habitat choice by both males and females can lead to size assortative mating even in the absence of an active mate choice. For example, smaller cichlid fish of both sexes prudently occupy vacant territories of worse quality than do larger fish, leading to size assortative mating [11]. Our results did not support this hypothesis. We found that smaller females used prudent burrow-site selection, but males exhibited similar sampling and burrow occupation behaviors, regardless of their body size and whether or not the sites were safe. This is probably because burrow-site selection by male isopods also integrates information about the quality of the resident females [17,27]. Thus, prudent burrow-site selection per se cannot lead to size assortative mating in our system.

Our findings imply that larger females should breed in safer sites leading to fear-phenotype matching across the landscape of fear. This expectation does not align with previous results from the end of the breeding season that show no differences in average female size between risky and safe sites [17]. Male isopods anticipate future costs of predation and use this information to lessen the expected reproductive benefit that is based solely on female size. Consequently, large males are less likely to settle in risky habitat leading to lower pairing rate in risky sites that may explain the much higher abandon rate of recently occupied burrows in risky sites compared to safer sites [17]. Higher burrow availability in risky sites at the end of the dispersal period may attract delayed female dispersers regardless of their size. This suggests that differential mate-choice by male isopods may interrupt the phenotype-habitat matching that is expected based on the prudent burrow site selection by small female isopods. Future studies should specifically test this plausible scenario.

In summary, we showed that larger female isopods were more likely to occupy new burrows, regardless of whether or not the burrow sites were safe. We also found that larger females stayed longer in safer sites while smaller female tended to stay longer in riskier sites, implying a prudent burrow selection by small female isopods. Male isopods show preference for larger females in safer sites raising the question of why smaller female despite the excessive fitness costs of not finding a mate or mating with a lower quality male still use prudent behavior. Future work should explore this conundrum by comparing the probabilities of smaller female to pair in risky and safer habitats. Our work highlighted the need to consider intraspecific competitions when exploring how predation risk affects prey breeding-site selection and mating patterns.

## Author contributions

Viraj R. Torsekar and Dror Hawlena conceived the study and designed the experiments. Viraj R. Torsekar and Aparna Lajmi collected the data and Viraj R. Torsekar analysed it. Viraj R. Torsekar and Dror Hawlena wrote the manuscript and Aparna Lajmi contributed to the final manuscript.

## Acknowledgements

We thank Nevo Sagi, Peleg Jakubovits, Yoann Pellen, Pnina Cohen, and Eyal Privman for help with conducting the field experiment, and the two reviewers and associate editor for providing valuable suggestion for improvent. The research was funded by European Research Council grant (ERC-2013-StG-337023 [ECOSTRESS]) and Israel Science Foundation grant (ISF-No. 1391/19) to Dror Hawlena.

## Ethics statement

Since we handled animals only briefly and released them meters away from the place of capture, we have no ethics approval to declare. Also, this species is not regulated by the HUJ ethics committee or protected by the Israeli law. Thus, no special permits were needed to conduct this research.

## Data availability statement

All the data analysed in this manuscript are available on DataDryad at https://datadryad.org/stash/share/7v6NVNEv1pY7znjjcbwQiHkK75ylAPEGdEK3_6-y0MQ Citation: Torsekar, Viraj; Lajmi, Aparna; Hawlena, Dror (2023), Prudent burrow-site selection in a landscape of fear, Dryad, Dataset, https://doi.org/10.5061/dryad.w6m905qvj

