## Supplementary information for "Prudent burrow-site selection in a landscape of fear"

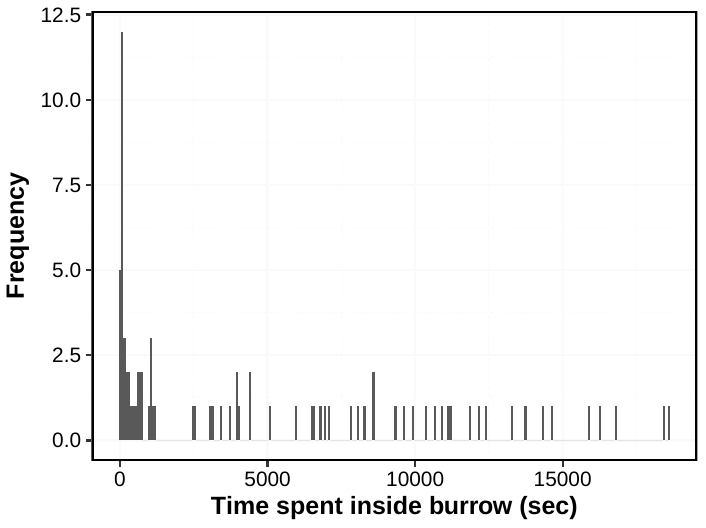

Fig. S1. Distribution of durations isopods spent inside every experimental burrow. The large gap in time durations between 1190 and 2400 seconds helped consider brief occupations (duration below 1190 seconds) as if the burrow is rejected.

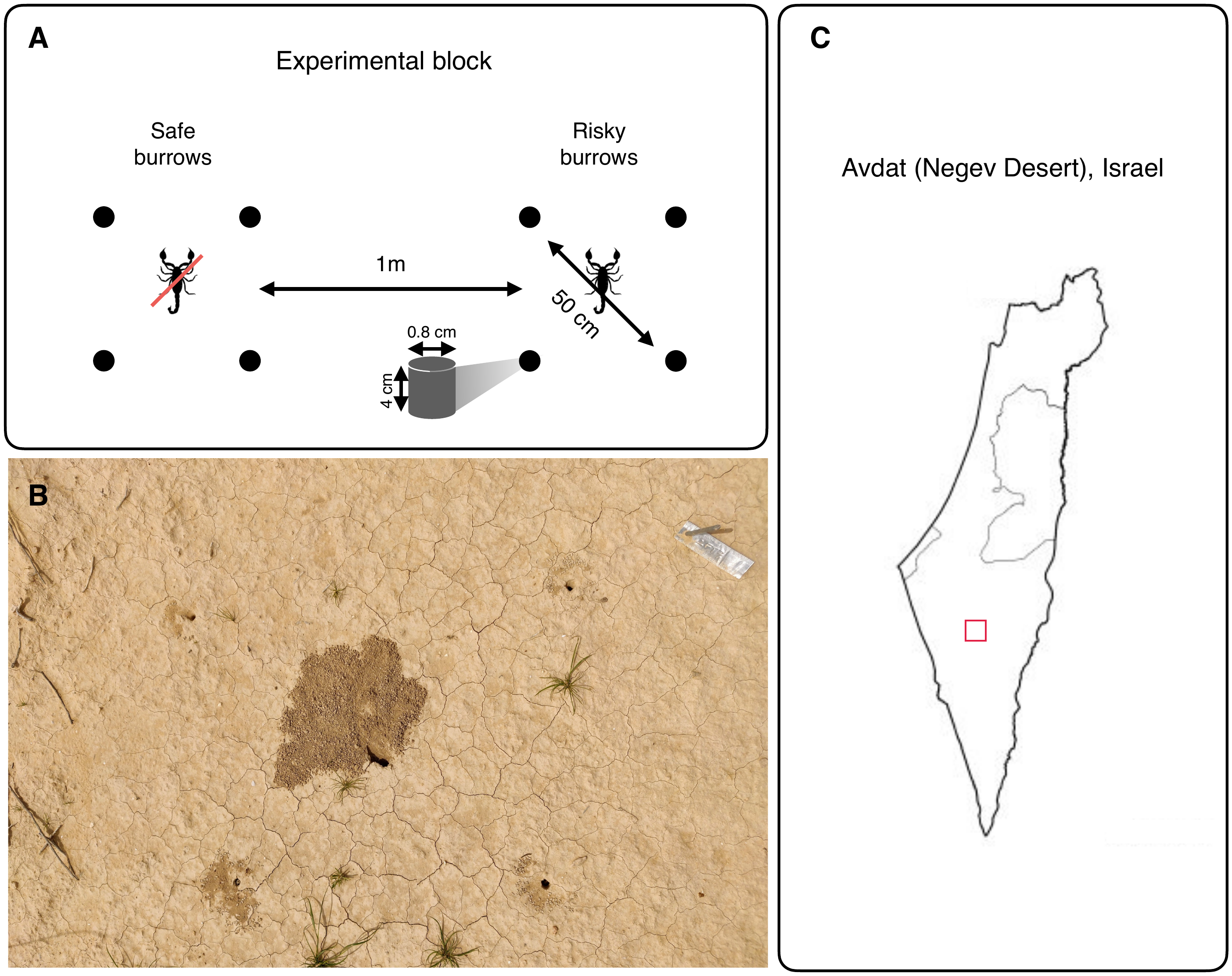

Fig. S2. Details of how and where the experiment was performed. A) Illustration of the experimental block. Each block included two groups of four 0.8 cm diameter x 4 cm depth cylindrical holes that were dug in the corners of an imaginary 50 cm diagonal square. The minimal distance between the two burrow-groups was one meter. In each block, we dug a 10 cm long scorpion-burrow in the middle of one randomly chosen burrow-group. The neighboring burrow-group in each block served as a no-scorpion “safe” control. B) Photograph of the risky burrow-group part of the experimental block with four dug holes and one scorpion burrow in the middle. C) Location of the Avdat Research Station, Negev desert, Israel (30°47’02” N, 34°46’09” E) marked using a red square.

**S1. Discounting of occupied burrows**

We removed isopod visits to already occupied holes from our analyses. These were a small proportion of all visits (22 out of 109) and they were similarly distributed between safe (N=7) and risky (N=15) burrows. Among the visiting isopods, 8 were females and 14 were males.
